# Aseptic, semi-sealed cranial chamber implants for chronic multi-channel neurochemical and electrophysiological neural recording in nonhuman primates

**DOI:** 10.1101/2025.02.10.636943

**Authors:** Jiwon Choi, Usamma Amjad, Raymond Murray, Ritesh Shrivastav, Tobias Teichert, Baldwin Goodell, David J. Schaeffer, Julia K. Oluoch, Helen N. Schwerdt

**Affiliations:** Department of Bioengineering, University of Pittsburgh, PA, USA; Aligning Science Across Parkinson’s (ASAP) Collaborative Research Network, Chevy Chase, MD, USA; Department of Psychiatry, University of Pittsburgh, PA, USA; Gray Matter Research, Bozeman, MT, USA; Department of Neurobiology, University of Pittsburgh, PA, USA

**Keywords:** primate cephalic chamber, chronic neural implants, neurochemical recording, multi-channel neural recording

## Abstract

We developed an implantable neural interface for monitoring both chemical and electrical forms of brain activity in monkeys that maintains aseptic properties for year-long periods while leveraging the modular functions (e.g., sensor moveability) provided by a chamber system. Invasive electrophysiological recordings, especially in subcortical structures from nonhuman primates usually involves implanting electrodes into the brain through a skull-mounted chamber. These electrodes may be attached temporarily for several hours of recording, or permanently. Permanent attachments are favorable to allow for sealing the chamber completely from externally originating pathogenic species that can infiltrate and compromise the health of the animal. A sealed chamber also reduces the need for frequent chamber cleaning required to minimize the accumulation of pathogenic organisms. However, neurochemical measurements require specialized electrodes with extremely fragile carbon fiber tips and are not compatible with recently developed sealed chamber systems. Here, we leveraged osseointegrating materials and hermetic sealing strategies to enable both neurochemical and electrical neural activity measurements from a sealed chamber with an aspirating port for culturing chamber fluid to ensure an aseptic environment. The system was shown to provide successful recordings of neural activity in two monkeys while maintaining negative bacteria culture results for over a year post-implant.

## 1. Introduction

Technologies for recording electrical and chemical neural activity in nonhuman primates have advanced tremendously over the last few decades^1–6^. Nevertheless, challenges remain in maintaining chronic (> months) measurements, and in achieving multi-site measurements in deep (> 5 mm), cortical and subcortical, brain regions. Methods that enable multi-modal, chemical and electrical, neural measurements are needed to elucidate the relationship between dynamic molecular (e.g., dopamine) and electrical neural activity (e.g., action potentials) changes known to be pivotal for learning and plasticity^7–9^. Such techniques could inform how these relationships are altered in pathological conditions such as in Parkinson’s disease. Expanding these types of multi-modal neurophysiological experiments in the primate species is also crucial from a translational perspective as it would allow us to pinpoint neural mechanisms and targets that have close homology to humans. We designed improved semi-sealed chamber systems to enable such multi-modal measurements over chronic timescales (> 1 year) in an aseptic interface in Rhesus monkeys.

Measurements of neural activity from deep brain regions of monkeys have been achieved predominantly using a “chamber” system^10–16^. The chamber is a sealable port fixed to the skull that can be opened for neurophysiological experiments to expose the brain surface. Chambers allow researchers to maintain access to the brain over several years, through which electrodes may be inserted to record or manipulate neural activity. Electrodes may be permanently implanted and attached to the chamber for chronic (also called “semi-chronic”) recording^4,10,11,16^. Chronic methods are highly advantageous as it increases the efficiency of neurophysiological studies by reducing the labor and time needed for preparing electrode insertion procedures in near-daily recording experiments. It also reduces the cumulative trauma induced as these recordings require piercing the brain with a guide tube needle used to protect the electrode during brain insertion and/or the electrode itself. Furthermore, chronic stability is necessary to study brain processes that evolve over the course of adaptation, learning, or disease processes, that evolve across days or years. Many of these chronic electrode harboring chambers are completely sealed interfaces, isolating the brain surface from the external environment. These systems obviate the need for labor intensive chamber-cleaning procedures to disinfect inside the chamber and minimize the accumulation of pathogenic microorganisms, which if left unchecked can cause significant issues, from bone resorption to meningitis^13^. This cleaning procedure usually must be performed 3 – 5 times a week, increasing the required labor and cost for maintenance. The implanted electrodes are protected by a cap to prevent damage when the monkey is freely roaming in the home-cage environment outside of experiments. Electrodes are mounted on micromanipulators or microdrives that allow chronically implanted electrodes to be lowered on demand, increasing the number of brain sites that can be sampled^4,10,11,16^. Such chamber systems have enabled a wealth of fundamental discoveries – from elucidating the diversity of functions harbored by specific brain regions^17^ to characterizing network level processes that occur across brain regions^18,19^.

To date, chamber systems have been mainly limited to chronic recording of electrical neural activity (e.g., LFPs and spikes) using standard electrophysiological (EPhys) methods. Recently, chamber systems were developed to enable chronic recording of both neurochemical activity using electrochemical, fast-scan cyclic voltammetry (FSCV), and electrical neural activity^1,2^. However, chronic neurochemical recording bears additional challenges in comparison to electrical recording techniques. Electrochemical techniques entail the use of delicate carbon fiber (CF) electrodes (CFEs) that are characterized by a small diameter (7 µm) tip and that are susceptible to fracture, with > 50% of these sensors experiencing mechanical failure at the tip during manual brain insertion procedures^2,4^. Successful insertion of these sensors has traditionally required the use of a guide tube, which the electrode is threaded through and that protects the tip from obstructions (e.g., dura mater) in the path towards the brain target (e.g., striatum, or caudate nucleus, CN, and putamen). Sealed chamber interfaces developed so far are not compatible with guide tube usage^10,16^ and therefore have not been attempted with CFEs. Here, we developed a chamber system that keeps the brain largely sealed from the external environment and thus reducing the risk of complications due to pathogenic species, while providing both chemical and electrical measurements with chronically implanted electrodes. Key advances made over prior chambers used for chronic neurochemical recording^1,2^ include a semi-sealed interface to restrict introduction of infectious species and remove the need for chamber cleaning, a sampling port to monitor chamber sterility, and the use of osseointegrating materials to further ensure a robust seal against the communication of environmental pathogens into the chamber, as well as a strong joint between the bone and the chamber.

## 2. Methods

### 2.1 Chamber Design

The chamber consists of three main components layered above one another: a baseplate, a top plate, and a cap (**Fig. 1**). The baseplate is attached to the cranium and is integral for: creating a robust attachment to the bone, isolating the brain surface from the external environment, and influencing the long-term health of the tissue and skin margins around the outside perimeters of the chamber as well as the tissue inside the chamber. The top plate attaches to the baseplate and heightens the overall chamber to create discretely sized cavities with enough space to accommodate fluid buildup from the brain surface, and to insert side windows for accessing the cavity. Finally, the cap covers the top plate and is used to protect subsequently implanted electrodes. Each component is detailed below.

**Figure 1.**
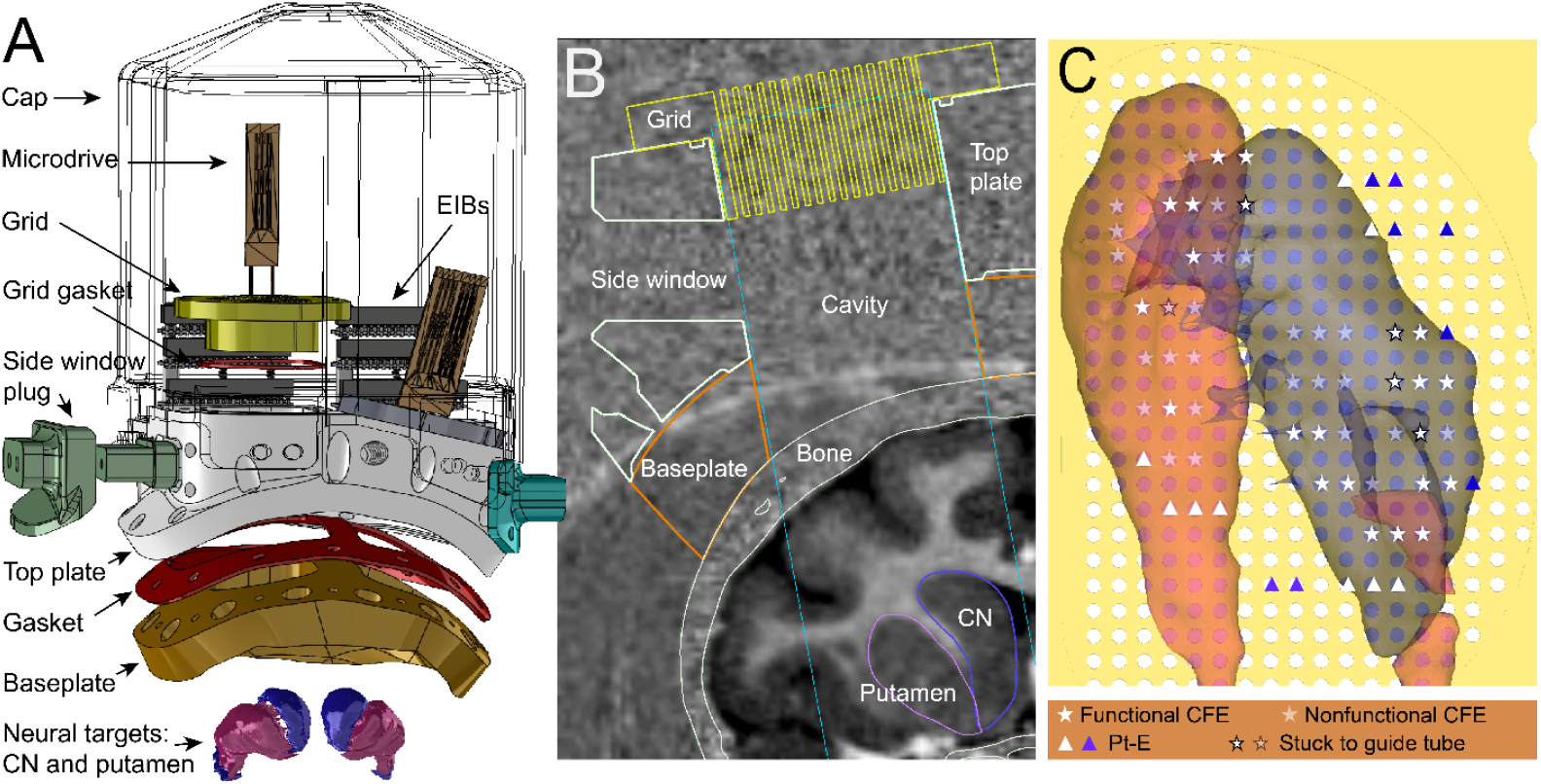
Overview of chamber design. **A.** Exploded view of chamber components (excluding implantable electrodes, aspirating port, screws for fastening, and window plug gaskets) and striatal targets (CN and putamen). **B**. Coronal view MRI co-registered with chamber components as would be implanted on monkey J. Boundaries of the skull, and the striatal targets are also outlined. **C**. View from above the grid, showing chamber encapsulated CN and putamen brain targets and electrode map showing grid holes in which CFEs or Pt-Es were implanted in monkey T and their fate for functional recording (see legend below).

**Figure 2.**
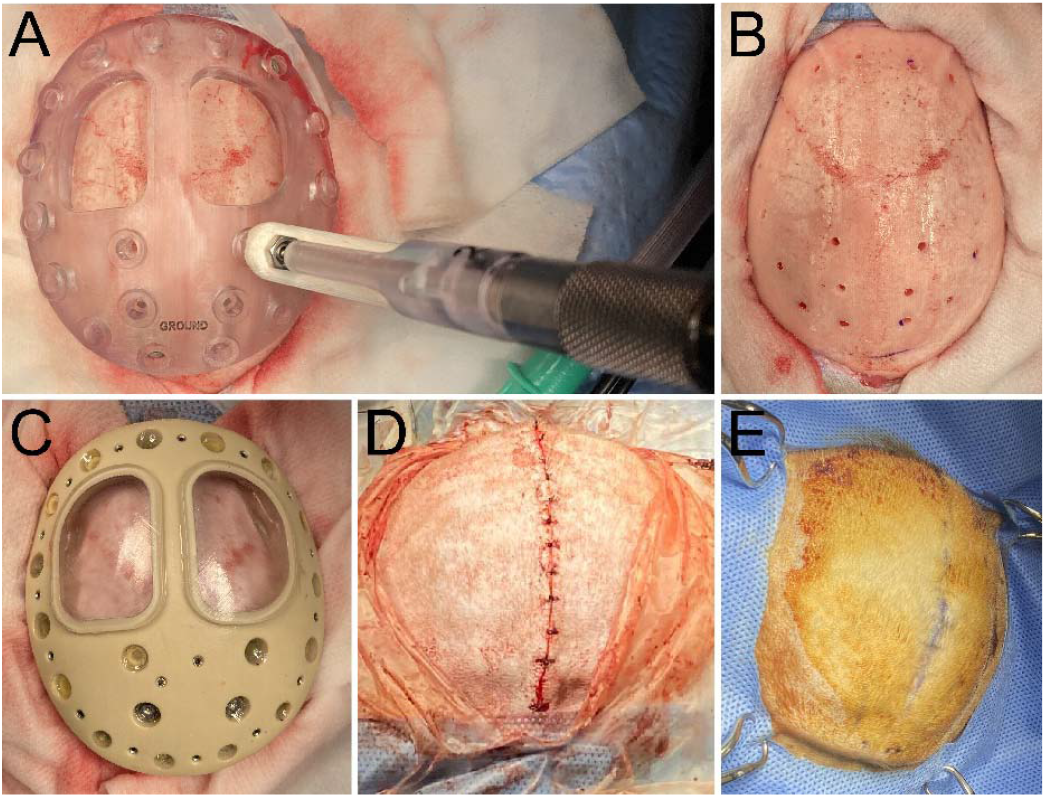
Baseplate implantation. **A.** Drill guide placed on skull for drilling intracranial screw holes into baseplate matching locations (monkey T). A drill sleeve was used with the drill guide to ensure that the hand drill produced holes at the minimal depths to allow complete penetration into the bone to prevent puncturing of the dura mater. **B**. Cranial surface after drilling all the screw holes with the drill guide (monkey T). **C**. Baseplate secured onto bone with titanium and ceramic screws (monkey T). **D**. Skin sutured to completely cover the baseplate (monkey J). **E**. 346 days after baseplate implantation in monkey J, and directly before the craniotomy procedure. The skin appears to be fully healed (blue markings on the skin are from a surgical pen used to plan the incision line for the subsequent procedure and the red color on the skin is from a betadine scrub used to disinfect the surgical field).

#### 2.1.1. Baseplate

The baseplate is an 8 mm thick structure made of 3D-printed polyetherketoneketone (PEKK), an FDA-approved biocompatible material that has been used for cranio-facial and spinal applications^20^. The baseplate anchors to the cranial surface with titanium or ceramic screws (as detailed in the “Implantation Procedures” section) (**Fig. 1A** and **B**). It has two “windows” or cavities that allow access to the cranial surface for subsequent craniotomy and electrode insertion procedures. A thin 3D-printed plastic lid was used to cover both windows and further sealed with a thin layer of polydimethylsiloxane (PDMS). The bottom surface of the baseplate was designed to conform to the geometry of the skull by applying computer-aided design (CAD) (Solidworks, RRID:SCR_024908) to the model of the bone extracted from the monkey’s CT scan. This conformal geometry allows us to directly attach the baseplate to the skull while ensuring a tight fit (i.e., minimal gaps between bone and baseplate) and without the need for acrylic cement. Most primate chambers are attached to the skull using polymethyl methacrylate (PMMA), also known as acrylic, dental, or bone cement. The cement may be applied in small coats or poured over the distributed bone-threaded screws to readily attach a chamber to the skull. However, these compounds frequently cause tissue damage, long-term bone resorption, unwanted granulation tissue growth and space in between bone and chamber implant which can also lead to communication of pathogens from the external environment. The surfaces of these materials are also known to be relatively susceptible bacterial adherence and biofilm formation^21^. Collectively, any combination of these issues may lead to failure over time, such as chamber detachment, and/or serious infection (e.g., meningitis). Thus, our chamber helps address these issues by removing the need for cement, in line with other recent advances in acrylic-free chamber designs^10,12–15^.

PEKK was chosen for use as the baseplate material given its enhanced osseointegration^22–24^ and anti-bacterial properties^20^, as compared to polyetheretherketone (PEEK), a similar material that comes from the same polymer family (i.e., polyaryletherketone, PAEK) and that has been used successfully for a wide range of biomedical implant applications^25^. This osseointegrating feature was targeted to enhance the integrity of the cranial attachment, by allowing bone to grow into the chamber orifices and around the cranial screws, thus reinforcing the interlock between the two interfaces. In addition, this bone growth is expected to help create a barrier against the bacteria that infiltrate and usually flourish at the chamber margins, and in the external environment^13^. Finally, the baseplate is thin enough (~ 8 mm) to be completely enclosed subcutaneously. Details of this procedure may be found in the subsequent “Implantation Procedures” section. This prevents external pathogens from being introduced inside the chamber, at the chamber margins, and in between the chamber and bone, during the process of recovery and osseointegration, further minimizing the risk of infection and reinforcing a healthy environment for the implant for long-term use.

#### 2.1.2. Top Plate

The top plate is a thicker layer (23 – 30 mm) that is screwed onto the baseplate (as detailed in the “Implantation Procedure’’ section) (**Fig. 1A** and **B**). The top plate is made of PEEK and has two vertical windows that align with the baseplate windows for allowing access to both hemispheres of the brain and for forming the cavity to accommodate a moderate buildup of fluid over time. It also consists of side ports that provide access to the inside of the chamber, if needed. These allow for cleaning or monitoring the inside of the chamber after electrodes are permanently installed when the top window would be sealed by the grid. The grid is made of 3D-printed plastic and consists of an array of holes used to insert and align microdrive-mounted electrodes with MRI guidance. The grid holes are spaced 1 mm apart from each other with a diameter of 0.47 mm (to fit standard 26 G needles), and extend over a range of 26 mm along the anterior-posterior (AP) axis, and 19 mm along the medio-lateral (ML) axis in the shape of an ellipse (**Fig. 1C**). Any of the grid holes can also be used to accommodate an aspiration port, where a tube is attached for channeling fluid in or out of the chamber cavity. These grids (Accura Clearvue material using a 3d Systems Projet 6000 HD printer) are secured on top of the vertical windows. Rubber gaskets are embedded in between mating surfaces that communicate with the inside of the chamber cavity, and the indwelling brain surface, to ensure that the brain remains sealed from the external environment. Gaskets were included in between the top plate and baseplate, the top plate and the grid, and the top plate and the side port plugs. Rubber gaskets are lined with high-vacuum grease (Dow Corning, 0131764) to reinforce the seal.

Threaded holes (#0-80) were made on the posterior side of the top plate to allow electrode interface boards (EIBs) to be screwed in for connecting implanted electrodes to external recording instrumentation (more details on this in “Preparation of Chamber and Implanted Devices”).

The exterior surface of the top plate (i.e., outside of the area that gets protected by the cap) contains sockets for 3 ball head screws. These ball heads are used for ball-and-socket connection to a head-fixation “halo” that connects to the monkey’s chair. Next to these ball heads are threaded holes for screws for the cap, described next.

#### 2.1.3. Cap

The cap is a dome-like cover that raises 80 – 100 mm above the top plate and encloses and protects all of the implanted electrodes, microdrives, and EIBs. This piece is made of an impact resistant 3D-printed plastic (Gray Matter Research, GMR) to ensure that it remains intact and securely protects internal components, when the monkey is roaming freely in the home-cage environment.

### 2.2. Preparation of Chamber and Implant Devices

Implanted electrodes consisted of carbon fiber electrodes (CFEs) for FSCV and EPhys recording^2,4^, platinum-iridium (PtIr) electrodes (Pt-E) (UEPLEDVMCN1X, FHC) for EPhys recording, stainless steel (SS) reference and ground wires, and Ag/AgCl reference and ground electrodes.

The fabrication of the CFEs are detailed elsewhere^2,4^ and consisted of both stiff and moveable silica-based CFEs (sCFEs), and micro-invasive CFEs (mCFEs). Electrodes were mounted onto microdrives that allow electrodes to be lowered in increments of 127 microns per screw turn, with a maximum of 15 mm travel^11^. Therefore, multiple sites may be sampled with a single electrode. Cyanoacrylate (Rhino Glue, Ultra) was used to attach electrodes to the microdrive shuttle. The relative length at which electrodes were affixed onto the microdrive shuttle was determined based on the targeted depth for each individual electrode and the targeted placement of the microdrive on the grid based on MRI mapping (see subsequent “MRI Guided Chamber and Electrode Placement”). All microdrive shuttles were elevated to the topmost position so that when the electrodes were inserted into the brain, they were at the most superficial depth. The targeting of the electrodes is discussed in the subsequent section (“MRI Guided Chamber and Electrode Placement”).

Reference and ground wires were created by cutting 30 – 50 mm long pieces from a spool of insulated SS and Ag (127 µm diameter, A-M Systems, 786000), exposing ~ 3 – 5 mm of the underlying metal on both sides using a razor to remove the PFA insulation, and then crimping a pin on one side (Mill-Max, 0489-0-15-01-11-02-04-0), for subsequent attachment to the EIB. The Ag/AgCl reference wire was created by soaking the exposed Ag on the uncrimped side in a solution of bleach (Clorox, Concentrated Regular Bleach), overnight. The wires would then be inserted superficially above the dura mater and/or into the brain. The insertion procedures are detailed in the “Implantation Procedures”.

Guide tubes were prepared to penetrate the stiff meningeal layers and allow electrodes to pass through and reach their brain targets without damage. 27 G SS needles (Connecticut Hypodermics, 27G XTW “A” Bevel) (0.292 mm minimum inner diameter and 0.4191 mm maximum outer diameter) were cut to lengths based on the desired insertion depth of the electrodes (see subsequent “MRI Guided Chamber and Electrode Placement”). This length (27– 40 mm) was calculated as the distance from the top of the grid to the targeted depth in the brain, plus an additional 2 – 3 mm of additional height. This additional height allowed it to be gripped, with a tweezer, above the grid, so that it could be raised out of the brain after the electrode had reached its targeted depth. Guide tubes were cut with a dicing machine and designed to reach 3 – 10 mm above the targeted electrode depth. The cut ends of these diced tubes were then cored with a lower gauge (18 – 23 G) beveled needle to maximize the opening of the tube, and lightly polished on a grinding stone to remove any sharp edges. Finally, the tubes were sonicated in acetone for 15 – 30 minutes (Fisherbrand, FB11203), and then a tungsten or SS rod was threaded inside the tube repeatedly to push out any remaining debris inside the tube in isopropyl alcohol.

EIBs were designed to conduct signals from implanted electrodes to external instrumentation (i.e., FSCV or EPhys recording amplifier headstages). The printed circuit board (PCB) has two sides of connectors with electrical traces drawn onto the board to join them. The side facing the chamber has multiple rows of female sockets (Mill-Max, 853-93-100-20-001000) that allow plugging of implanted electrodes. The distal side has a standard small multi-pin connector (Omnetics, A79026-001, A79024-001) that allows plugging of FSCV or EPhys headstages.

The aspirating port consisted of a standard 26 or 23 G SS needle connected to flexible PE/PVC tubing (Instech, BTCOEX-25) that was then plugged in with a SS rod (oversized 0.02” diameter and 7/8” long) (McMaster Carr, 3023A394). The collection needle was placed inside the grid with the tip 5 – 10 mm above the brain surface to allow aspiration of fluid within the chamber without being occluded by subsequent growth of granulation tissue.

All components were sterilized in ethylene oxide gas before the electrode implantation procedure. Prior to the electrode insertion procedure, sterilized guide tubes were filled with sterile petrolatum lubricant (Dechra, Puralube Vet Ointment) and then inserted with the microdrive-mounted electrodes in a sterile environment. This gel helps create a seal in the guide tube to prevent bacteria and other infectious species from communicating into the brain through the guide tube. The viscosity of the ointment helps suppress back-flow of extracellular fluid or blood out of the tube during the insertion process. If this happens, the fluid will evaporate over time and leave behind salt crystals and other physical residue that can create a permanent, glue-like, attachment between the electrode and the guide tube, limiting any subsequent movement or lowering of the electrode. This gel also helps keep the electrode lubricated for subsequent lowering. These guide tube-electrode assemblies were then submitted for gas sterilization a second time.

### 2.3. MRI Guided Chamber and Electrode Placement

The location of the chamber and electrode trajectories were planned using 3D Slicer (RRID:SCR_005619) and based on the brain regions targeted for recording (i.e., CN and putamen) using MRI images co-registered with the CT scans used for building the baseplate surface (**Fig. 1B** and **C**). CT scans (250 μm isotropic) were acquired in an Epica Vimago GT30 animal scanner. T1-weighted MRIs were acquired for presurgical planning at the University of Pittsburgh on a human, whole body 7T MRI scanner (MAGNETOM, SIEMENS, Germany) using a MP2RAGE (magnetization-prepared 2 rapid acquisition gradient echo) sequence and a 2nd generation Tic-Tac-Toe radiofrequency (RF) system (resolution: 400 μm isotropic, TR/TE: 6000/3.7 ms, FOV: 148×148×96 mm), with 60 transmit channels and 32 receive channels^26,27^. The area of the chamber windows were designed to maximize coverage of the striatum on each hemisphere. We also adjusted the insertion angle to be 0 – 14 degrees off of the coronal plane. An angled approach allowed targeting of medial CN while avoiding passing electrodes through the overlying lateral ventricle. The 100 – 300 µm long CF tips of the CFEs have been found to break during their passage from the ventricle to cerebral tissue, most likely due to the rigidity of the arachnoid-pia membrane that encases the brain. Electrode insertions were simulated in 3D Slicer with models of the chamber and grid integrated with the MRI scans to determine trajectory lengths and coordinates for implanted electrodes. Electrodes could be implanted as closely as 1 mm apart (i.e., grid-hole spacing). Electrodes were carefully mounted onto a microdrive, which would also determine the reach of the electrodes in the brain. Up to 6 electrodes could be installed on a single microdrive. Electrode trajectories were simulated in 3D Slicer such that the tips of the electrodes reached just 1 – 2 mm dorsal to the surface of the striatum, allowing us to lower these more carefully through screw-drive manipulation of the microdrive shuttle (1 turn ~ 158 µm). This subsequent lowering into the targeted brain region occurred several months after the initial electrode insertion to allow recovery from the electrode insertion-induced trauma and tissue responses.

A total of 49 – 59 CFEs were targeted for the striatum (18 sCFEs and 8 mCFEs in the CN and 22 sCFEs and 6 mCFEs in the putamen for monkey T; 36 sCFEs and 8 mCFEs in the CN and 15 sCFEs and 6 mCFEs in the putamen for monkey J) for FSCV and EPhys recording, 17 or 20 Pt-E’s in monkey T and J, respectively, were targeted for cortical (i.e., primary motor cortex and frontal eye field) and subcortical regions for EPhys recording, and 3 – 5 Ag/AgCl wires were inserted into various white matter locations distal from the targeted recording sites to serve as references for FSCV recording. 5 – 10 Ag/AgCl and SS wires were to be placed on the surface of the dura mater.

### 2.4. Implantation Procedures

The surgical implantation procedures were divided into three main phases, each done on separate days: baseplate implantation, top plate attachment and craniotomy, and electrode implantation. All procedures were performed on 2 Rhesus monkeys (T and J) and were approved by the Institute’s Animal Care and Use Committee (IACUC) at the University of Pittsburgh and were performed following the Guide for the Care and Use of Laboratory Animals (Department of Health and Human Services), the provisions of the Animals Welfare Act (USDA) and all applicable federal and state laws. All implanted devices were sterilized in ethylene oxide gas or hydrogen peroxide plasma chambers. All procedures were performed in a sterile environment, and with the approved use of anesthetics, anti-inflammatory, analgesic, and prophylactic antibiotics.

#### 2.4.1 Baseplate Implantation

Monkeys were first given ketamine and atropine in their home-cage and then maintained on anesthesia with 1.5 – 2.0% isoflurane and 1 L/min oxygen. Analgesics, anti-inflammatory agents, and prophylactic antibiotics were administered pre-and/or post-op (i.e., meloxicam, dexamethasone, ceftriaxone, and buprenorphine). Monkeys were placed in stereotactic frames to fix their head for surgical operation. A sterile field was created on the skin above the targeted location for the baseplate and the surface was disinfected with several rounds of applying betadine and 70% isopropanol, in a serial manner, with gauze. This same monkey preparation procedure was performed at the start of all subsequent procedures described.

The cranial surface was exposed by making an incision along the midline of the scalp. The baseplate was then fit onto the skull. Proper placement of the baseplate was important to match the pre-planned MRI orientation and ensure accurate targeting of subsequent electrodes. This was done by moving the baseplate around the surface of the skull until a position was found that restricted significant lateral movement meaning that it had conformed to the bone geometry (i.e., the baseplate had a tight fit on the surface). A sterile pen was used to mark this placement by drawing an outline on the skull around the baseplate. A drill guide plate having the same shape as the baseplate was used to first drill holes into the bone for subsequent threading of intracranial screws to secure the baseplate to the bone. The drill guide was placed in the same location as the baseplate, based on the outline that was drawn on the skull and the fit. This drill guide was used to restrict the drill’s drill-bit to a depth equal to the thickness of the skull (~ 2 – 3.5 mm) and to prevent the drill from piercing the dura mater. The drill guide had 20 clearance holes that were aligned to the screw hole locations on the baseplate. These holes were sized to closely fit the drill-bit (2 mm diameter). The depth of the hole produced in the bone was controlled by a drill-sleeve that controlled the depth with 0.5 mm resolution.

After intracranial holes were drilled, the drill guide was removed and the baseplate was placed back onto the bone and aligned to the holes. If needed, manual drilling was performed to widen holes that were misaligned between the two plates. Significant misalignment occurred in 5 (monkey T) or 2 (monkey J) holes. The baseplate was then screwed onto the skull using ceramic and titanium screws (2.7 mm diameter, from Gray Matter Research, GMR). Custom-made SS ground screws (GMR) were also fastened into two locations, to be used for providing electrical ground connections to the animal skulls. The screws were then covered with a fast-curing silicone (WPI, Kwik-Sil), except for SS ground screws. Ground screws were covered with a thin plastic cover and a drop of Kwik-Sil on top to protect the underlying electrical socket from debris and to keep it clean for subsequent electrical connection. The retracted skin was then pulled over the baseplate and sutured using two-layers of 3-0 sutures (dissolvable suture beneath the skin and non-absorbable nylon suture for over the skin) with simple interrupted patterns. The details of this procedure are also available at protocols.io (dx.doi.org/10.17504/protocols.io.kqdg32b91v25/v1).

#### 2.4.2. Craniotomy and Top Plate Attachment

The same procedures to prepare the monkey for sterile surgery were followed as in the “Baseplate Implantation” procedure described previously. The skin was incised down the center of the baseplate and retracted to expose the baseplate surface.

At this point, either an intermediate top plate was installed (monkey T), or a craniotomy was performed along with the final top plate installation (monkey J). In monkey T, an intermediate protective plate was installed because an open wound at the anterior and posterior edge of the baseplate had developed 1.5 months after the baseplate had been implanted. Thus, we decided to open the incision to clean and treat the wound and to prevent bacteria from accumulating inside the wound and around the baseplate. The exposed baseplate was cleaned thoroughly with diluted betadine and rinsed with saline. The protective plate was then screwed onto the baseplate to protect the baseplate surface and to keep the cranial window covers sealed to prevent the underlying bone from being exposed to the external environment. Sutures were used to close any exposed bone and create a tight seal between the skin and the chamber margin. Thus, the craniotomy was performed in a separate (fourth) procedure for monkey T (304 days after the initial baseplate implantation). Monkey J represented the ideal case, proceeding directly to craniotomy and final top plate installation in the second procedure (346 days after the initial baseplate procedure).

Craniotomies were performed by first removing the baseplate window cover to expose the bone surface (**Fig. 3A**). A foot-pedal-operated electric drill with a stainless steel or diamond-coated burr bit was used to thin down the bone surface and create a hole to expose the brain tissue. Throughout the process, 0.9% saline was perfused on the bone and exposed brain to prevent over-heating from the electric drill as well as to keep the tissue moist. A rongeur was also used to clip away the remaining bone inside the window. A bone curette was then used to scrape away any protruding bone remaining at the edges of the cranial window. In both monkeys, craniotomies were only performed on the right hemisphere window. The left hemisphere window was left intact to be used for future procedures.

**Figure 3.**
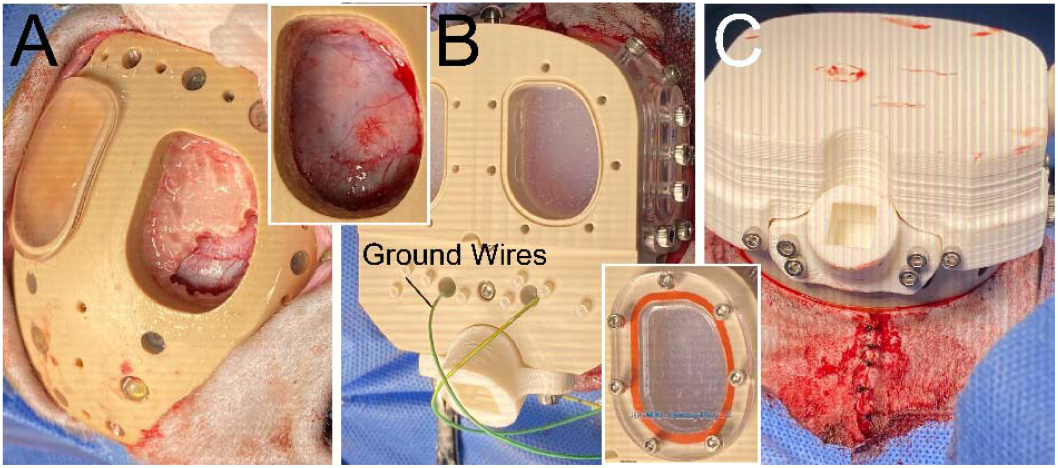
Craniotomy and top plate attachment (monkey J). **A.** Baseplate exposed beneath the skin, and bone is carefully removed using a motorized drill and rongeurs for the right hemisphere port. Top right inset shows the complete removal of the bone and exposed dura mater. **B**. Top plate is screwed onto the baseplate, the chamber cavities are filled with a fast-cure silicone (Kwik-Sil), and ground wires are installed onto the ground screws. Bottom right inset shows the cover installed for the chamber cavity, which will be replaced by a grid in subsequent procedures. **C**. Cap installed onto the top plate and the skin margins are sutured around the assembled chamber to ensure that bone is not exposed.

The final top plate was installed after the craniotomy for monkey J, and 2 months after the intermediate protective plate in monkey T (before the craniotomy) (**Fig. 3B**). Performing the craniotomy before final plate installation helps allow easier access to the bone with the drill because of the shallow height of the baseplate by itself. Ground wires were first plugged into the two grounding bone-screws in the baseplate and threaded individually into channels running through the top plate. The top plate was then screwed onto the baseplate with an interleaving rubber gasket used to seal the interface between the two pieces. The rubber gasket was lightly covered with high-vacuum grease (Dow Corning, 0131764) to help reinforce the seal. Both cranial windows were filled with Kwik-Sil to seal the exposed brain (right hemisphere) or the bone surface (left hemisphere). A cover was then installed onto both windows, with a rubber gasket, to further reinforce the seal and ensure a robust aseptic enclosure. A protective cap was screwed onto the top plate. In monkey J, the skin was sutured as done in monkey T for the intermediate top plate procedure (**Fig. 3C**). The details of this procedure are also available at protocols.io (dx.doi.org/10.17504/protocols.io.x54v92wd4l3e/v1).

#### 2.4.3. Electrode Implantation

The protective cap was removed to expose the top of the chamber and its windows. The top of the chamber was disinfected by rubbing 70% isopropanol infused gauze across the surface several times. The surrounding chamber and implant margins were disinfected by wiping with gauze containing diluted betadine and/or, separately, 70% isopropanol, in a sequential manner. Care was taken to avoid applying 70% isopropanol at the margins. A sterile field was created by isolating the top of the chamber with sterile drapes.

Electrode interface boards (EIBs) were covered with a transparent silicone coating (MG Chemicals, 422C) and were installed onto the chamber with screws that threaded into the top plate (**Fig. 4**). The bone-screw ground wires that had been exposed in the previous procedure (see “Craniotomy and Top Plate Attachment”) were plugged into the EIB ground input connectors. The rest of the EIB connectors would be used to connect subsequently implanted electrodes. The window cover for the right hemisphere (i.e., the window in which craniotomy was previously performed) was removed from the chamber, along with the indwelling Kwik-Sil plug. This exposed the brain surface (i.e., the surface of the dura mater and overlying granulation tissue). Only one hemisphere was targeted for this procedure, and the window targeting the opposite hemisphere remained closed, for use in future procedures. The MRI-mapped grid was installed to replace the previously installed window cover on top of the window and also with a rubber gasket and vacuum grease to reinforce the seal. On this grid, the targeted holes for inserting electrodes were marked with a black permanent marker to enhance visibility and facilitate manual alignment of electrode assemblies through these holes. All other holes on the grid were filled with caulk (DAP, Kwik Seal Ultra, 49395), except for the holes in which microdrive posts were to insert and attach into. The side window cover was also removed to allow for aspirating of liquid build up resulting from electrode insertions, as well as to rinse 0.9% saline into the chamber when bleeding occurred. This also allowed us to visualize and confirm that the trajectories of inserted electrodes were perpendicular relative to the grid surface to reduce positioning errors.

**Figure 4.**
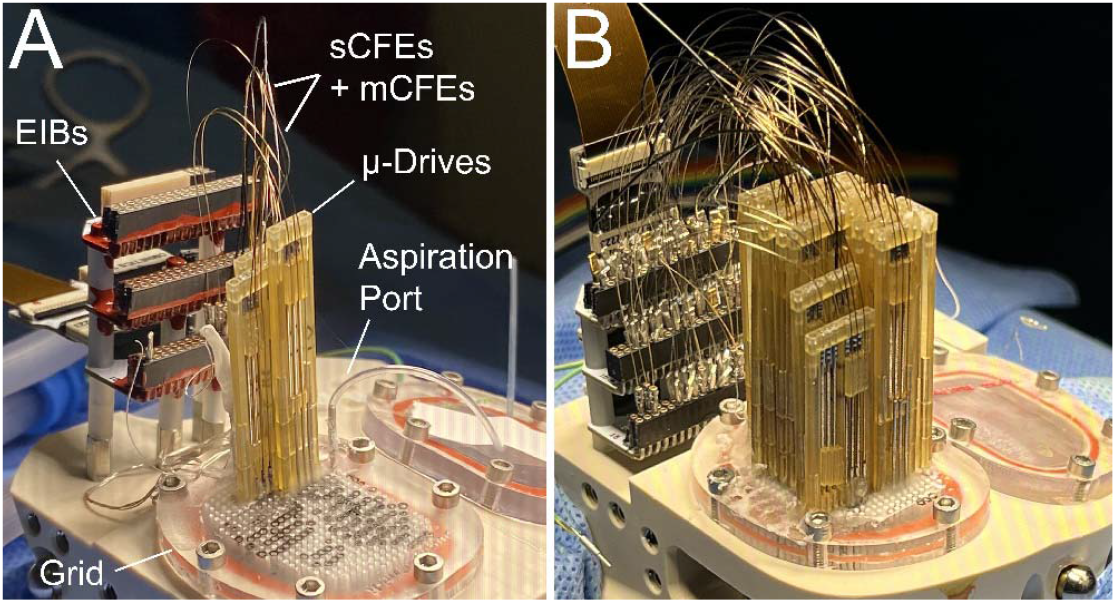
Electrode implantation. **A.** In the process of electrode implantation (monkey J), chamber grid has 3 mounted microdrives, and the aspiration port. The implanted CFEs and reference wires are connected to the EIB mounted in the posterior of the chamber. The grid is marked for placement of subsequent electrodes. **B**. Fully implanted chamber with 17 microdrives (monkey T).

Multiple (3 – 5) superficial Ag/AgCl and (3 – 5) SS reference wires were inserted into the grid. These were positioned so that the exposed portions laid flat on the surface of the brain while remaining in the posterior edge of the chamber and outside of the electrode insertion fields. 3 – 5 intracranial Ag/AgCl wires were also installed to target the white matter area of the brain, and far away from brain structures targeted for neural recording. In this case, a 27 G needle was used to gently penetrate the surface of the brain to create an opening in the dura mater. This was followed by inserting a 27G guide tube pre-threaded with the Ag/AgCl wire into the targeted area in the white matter. The Ag/AgCl wire was then pushed in deeper through the guide tube so that it’s tip extended 3 – 5 mm from the guide tube tip so that the entire exposed portion of the wire was inside the brain.

Microdrive-mounted electrodes were inserted into the brain starting with the most posterior placements and each additional drive being placed more anterior (**Fig. 4A**). The targeted grid holes for the electrodes mounted on a given microdrive were penetrated with a 27 G needle to remove any caulk or debris residing in the grid hole. A 27 G needle was then used to pierce the surface of the brain and create an opening in the dura mater. This process was done before installing each electrode / microdrive assembly to the chamber. Two surgeons worked together to insert the electrodes carefully in the brain while ensuring that the microdrive was lowered slowly and in the correct direction. All electrodes were coated with sterile petrolatum lubricant (Dechra, Puralube Vet Ointment), which was also filled in the guide-tube, as described in “Preparation of Chamber and Implant Devices”. The first surgeon positioned the microdrive with attached electrodes on top of the targeted holes above the grid. Guide tubes enclosing the hanging electrodes did not slip off the electrodes due to the adhesive forces provided by the eye-ointment gel. The second surgeon held the microdrive with tweezers. This was done to prevent the microdrive from falling or moving too fast as the first surgeon lowered it manually onto the grid. During the microdrive lowering, the guide tubes were first lowered and penetrated the brain. The electrodes remained inside the guide tubes and their tips were retracted from the tips of the guide tubes to keep them protected during the dura mater penetration. Once all the guide tubes had advanced into the brain, the microdrive assembly continued to be lowered, and the electrodes and guide tubes together were lowered into the brain. Guide tubes were no longer lowered once reaching their targeted depth, usually 5-10 mm above the electrode target. The guide tubes were cut to discrete lengths based on the electrode’s targeted depth and such that at least 2 – 3 mm of the guide tube remained above the grid. At this point, only the electrodes and microdrive were lowered together until the microdrive reached the surface of the grid and the electrodes reached their targeted depth.

Guide tubes were slowly lifted out of the brain, and placed just above the brain surface, after the microdrive was securely attached to the grid. Surgical tape (McKesson, 19-0772) was placed on the grid just in front of the guide tubes. This tape acted as a protective mask to prevent the liquid acrylic cement (Lang Dental, Jet Denture Repair Powder and Liquid), that would be used for the microdrives, from entering and covering the more anterior holes. A paper point tip (EMS Diasum, 71011-01) or applicator spear was used to apply a small amount of acetone on the guide tubes and the surrounding grid surface to remove any eye-ointment residue, which would compromise the adhesive strength of subsequent acrylic cement. Cement was applied in the grid holes surrounding the guide tubes, between the guide tubes and the microdrive, and at the perimeter of the microdrive where it attached to the grid to further reinforce attachment of the microdrive to the grid. The masking tape was then removed when the cement had hardened to a gel like consistency, but before it had fully cured. Electrodes were then connected to the EIB, which was then connected to the FSCV amplifier headstage to allow recording of signals from the electrodes. Working electrodes were noted based on visualizing electrochemical current > 500 nA. Electrodes displaying current less than this threshold were noted as broken. The chamber cavity was aspirated and rinsed with 0.9% saline through the side port. This electrode lowering process was repeated for each microdrive starting from the posterior edge of the chamber and adding microdrives one after another in the anterior direction.

After all microdrives had been installed (**Fig. 4B**), red varnish (GC Electronics, 10-9002-A) followed by white liquid insulating tape (Blue Magic, BOT59TRI), was applied onto the Mill-Max connectors where electrodes were plugged into the EIB boards. This was done to maintain a physically robust connection and prevent future animal movements and vibrations from weakening the junction between the electrodes and the EIB connectors, which is a common failure mechanism^4^.

An aspiration tube was installed onto the grid either during this procedure (monkey J) or 3 days after (monkey T). The aspiration tube consisted of a 23G (monkey J) or 26G (monkey T) needle attached to flexible plastic tubing (10-15 cm long) using cyanoacrylate glue (Rhino glue). The needle was inserted into the grid and several millimeters above the brain surface and then secured and sealed to the grid using Kwik-Sil. Kwik-Sil was used instead of a stronger adhesive (e.g., acrylic cement or cyanoacrylate) to allow the needle to be easily removed in the case that it became obstructed and had to be replaced in the future. The tube was closed with a SS plug (McMaster-Carr, 0.02” diameter, 7/8” long, hardened oversized high-speed M2 tool steel rod) at the other end.

The final steps involved aspirating any remaining fluid inside the chamber from the side window, closing the side window, and then placing the tall protective cap to enclose all the implanted electrodes and microdrives above the chamber. The details of this procedure are also available at protocols.io (dx.doi.org/10.17504/protocols.io.bp2l62m95gqe/v1).

### 2.5. Post-Operative Maintenance

Aspiration of chamber fluid was performed to prevent excessive fluid buildup in the chamber cavity, which could eventually leak through the guide tubes and out of the chamber (**Fig. 5**). In previous studies^2,4,11^, this was done by opening side windows to access the chamber cavity. However, the exposure provided by the relatively large opening (~ 5 – 10 mm diameter) of the windows, and the weak seal between windows and their covers would lead to eventual infiltration of bacteria within the chamber. This is a common issue in most chambers, including those used for acute recording experiments^13,14^. Due to this contamination, near-daily aspiration and cleaning (e.g., flushing with diluted betadine and 0.9% saline) within the chamber is required to suppress significant growth and spreading of the bacteria. Omission of such cleaning procedures usually leads to meningitis or other fatal infections. The use of water-tight seals such as rubber gaskets in our chamber design and only exposing a small channel for aspiration with an aspiration tube likely reduced the risks of bacterial contamination.

**Figure 5.**
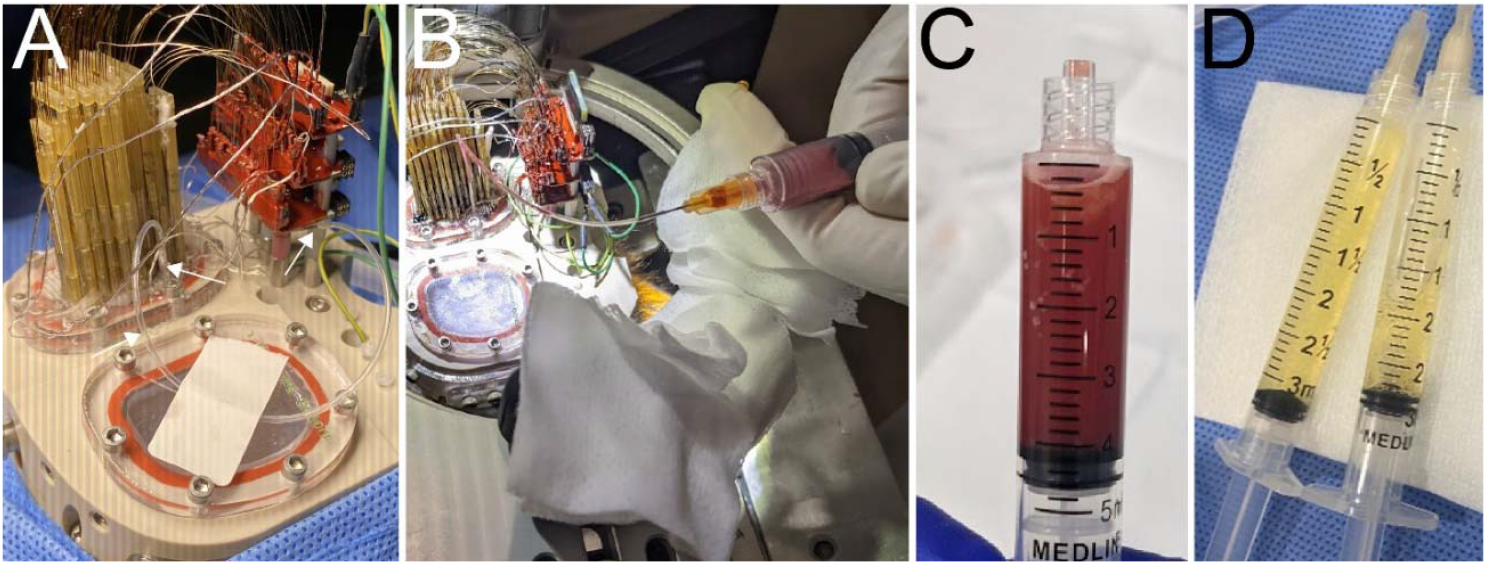
Aspiration process. **A.** Aspirating port and tube are shown in monkey J. Arrowheads indicate the aspirating port and the tube plug. **B**. Syringe is connected to the tube and used to manually pull liquid out of the chamber activity. **C**. Example sanguinous fluid collected. **D**. Example of clear serous fluid collected.

We aspirated fluid through the aspiration tube that was installed during the final electrode implantation procedure. All aspirations were performed in an aseptic environment with the monkey chaired and with the head fixed and anchored to the chair using a “halo” system (Gray Matter Research). The cap was unscrewed from the chamber to reveal the enclosed electrode implants and aspiration tube. The aspiration tube was disinfected around the opening by rubbing with 70% isopropanol. The SS rod was unplugged from the tube and a 26 G (monkey T), or 23 G (monkey J) needle connected to an empty syringe (usually 3 – 10 mL capacity) was inserted into the tube. The plunger was pulled outwards on the syringe to generate sufficient negative pressure to withdraw fluid from the chamber. In some cases, a larger syringe (e.g., 20 mL) was used to generate more negative pressure when the fluid was more viscous (e.g., coagulated blood). This withdrawal took place until only air was withdrawn. A small amount of the withdrawn fluid was submitted for processing identification and susceptibility for aerobic and anaerobic cultures (IDEXX BioAnalytics). Aspiration was performed near-daily immediately after the surgery, when we noticed a significant rate of fluid buildup. Once the rate of fluid buildup decreased, the aspiration was performed less frequently, and after a few months, aspiration only needed to be performed monthly.

### 2.6. Recording Chemical and Electrical Neural Activity

The procedure for recording chemical and electrical neural activity is detailed in existing work^1,2^. Briefly, dopamine neurochemical signaling was recorded using fast-scan cyclic voltammetry (FSCV) and electrical neural activity using standard electrophysiology (Neuralynx). A subset of sensors were connected to the FSCV system for recording dopamine concentration changes (Δ[DA]) and the remainder were connected to the electrophysiology system for recording spikes and LFPs. FSCV measures current generated by reduction and oxidation (redox) of targeted molecules, such as dopamine, by applying a ramping voltage from –0.4 to 1.3 V at the implanted CFE. Redox occurs at around –0.2 and 0.6 V for dopamine. These voltage-dependent current changes can then be converted to the estimated Δ[DA] using PCA and linear regression^2,28^. Standards of dopamine, pH, and movement artifact are used to further isolate selective current contributions associated with dopamine.

Measurements were made as monkeys performed a simple reward-biased eye-movement task, previously described^2,4^. The task involved a series of trials. A trial started with a central fixation cue that monkeys had to fixate for 1 – 2 s, after which a peripheral target was displayed that monkeys had to saccade to and fixate for another 2 – 3 s. After these series of saccades and fixations were made successfully, a feedback cue was displayed (a blue outline around the peripheral target cue) and a large or small liquid reward (diluted apple juice) was delivered. The size of the reward depended on the location of the target cue (top or bottom of screen). This task allowed us to measure and identify neural signaling changes related to reward.

## 3. Results and Discussion

### 3.1. Recovery from Procedures

For most of the invasive procedures, both monkeys displayed normal recovery without overt effects. However, in monkey J, we observed weakening and decreased use of the left hand and arm, contralateral to the implanted hemisphere, and lethargic behavior 4 days after the electrode implantation procedure. We increased the dose of anti-inflammatory drugs (i.e., dexamethasone) given to 1 mg/kg, as well as the administration frequency (i.e., from once to twice a day), for the next week (the monkey was on a lower, daily, 0.3 mg/kg dose for 4 days post-procedure). This dose was gradually tapered down to 0.125 mg/kg for the final 3 days of treatment. Symptoms gradually reduced and the monkey appeared normal in behavior after this period. We suspect that the hemiparesis may be due to trauma induced by densely packed guide tubes that penetrated motor pathways in the cortex and underlying white matter. The guide tubes are significantly larger (~ 0.4 mm diameter) than the implanted electrode and more than 82 of these were used within a small area (the area of brain surface exposed within the inner chamber was 519 mm^2^ for monkey J), which may have caused excessive brain damage due to the cumulative size of the implant.

### 3.2. Chamber Fluid Drainage

Fluid collected from the chamber aspiration tube for a given day varied between 0.3 – 4.8 mL for monkey T and between 0.2 – 9.8 mL for monkey J (**Fig. 6**). The estimated volume of the chamber cavity, with the bone removed, was 9.39 cm^3^ in monkey T and 10.16 cm^3^ in monkey J based on CAD designs. We aimed to drain the chamber fluid at frequent enough intervals to prevent fluid buildup beyond chamber capacity, and this frequency was largely based on the estimated perfusion rate from volumes collected in previous sessions (**Fig. 6B**). The sample collected from the aspiration tube was sanguinous (i.e., bloody) in the first few weeks or months (18 days post-implant in monkey T and 217 days in J) (**Fig. 5C**). In some cases, the blood coagulated and became too viscous and/or obstructed tube drainage completely. In these cases, the aspiration needle-tube assembly was carefully replaced after disinfecting the area around the grid, and aspirating manually through the hole in the grid (without a tube to reduce fluid resistance). After this period, we noticed that the fluid collected was clear, in the form of serous fluid (**Fig. 5D**).

**Figure 6.**
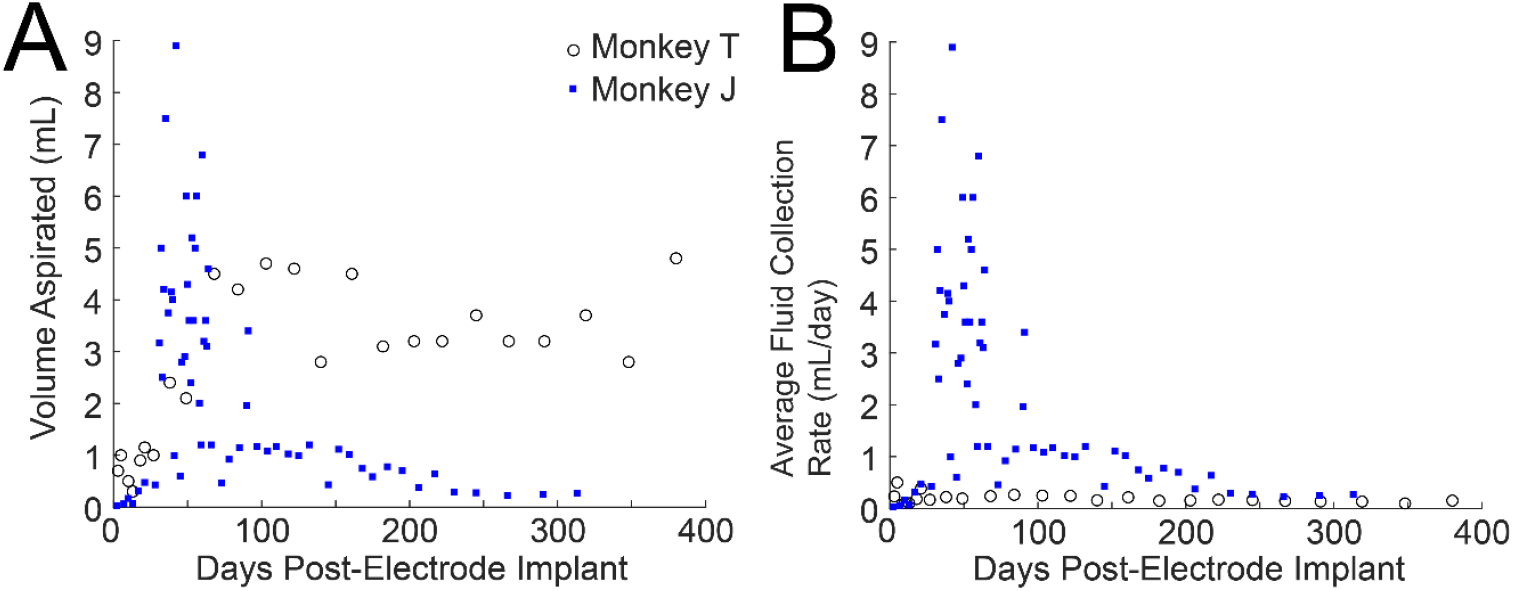
Drainage volume from chamber cavities. **A.** The volume of fluid collected from the chamber cavity during each aspiration session is plotted as a function of days post-electrode implant. Note that the sampling interval is shorter in general for monkey J and therefore less fluid may have accumulated in the later sessions (> 100 days) in comparison to monkey T. **B**. The average rate in which fluid was collected is plotted to normalize for the differing intervals between aspiration between monkeys and on a session-to-session basis.

In monkey J, the fluid alternated between serous and sanguinous forms over multiple drainage sessions, which contrasted with monkey T, where the fluid remained consistently transparent and without obvious blood after 18 days post-implant. We suspect that the sanguinous discharge may be a result of more highly vascularized granulation tissue forming above the dura mater and/or its interaction with the implanted devices in this monkey. Tube obstructions were addressed without needle replacement after several repeated cases of obstructions were observed in monkey J. Instead, saline was pushed through the tube into the chamber to reduce the fluid’s viscosity and then in most cases, we were able to withdraw liquid (both the expected discharge from the chamber, as well as the perfused volume) from the chamber successfully. This helped to reduce repeated removal and re-installation of the aspiration needle, which could lead to eventual exposure and compromising the sterility of the chamber.

Bacteria (staphylococcus) was cultured about a month post-implant (29 days) in monkey J, which we were able to successfully treat and remove. Monkey T maintained negative cultures for all collected samples. We suspect that the contamination in the chamber cavity for monkey J came from transmission through the guide tubes or surrounding grid holes as bloody fluid was observed in these areas on several occasions. Acrylic cement was applied in the areas where we suspected leakage, after drying the area with cotton swabs and applying acetone. 4 days following this result and the observation of repeated presence of bacteria, we began infusions of diluted antibiotics into the chamber. Infusions consisted of 1 mL of 3 – 16.4% Cefazolin in 0.9% saline, except on day 32, 33, and 39 post-implant when the infusion volume was 0.8, 1.5, and 1.5 mL, respectively. Each infusion took place after aspirating chamber fluid, and was left overnight. During this time, aspirations took place near daily and the concentration of the antibiotic was increased over time to prevent generating antibiotic-resistant bacteria. Also, the monkey was maintained on systemic antibiotics (Ceftriaxone). 39 – 44 days post-implant the samples began to show negative results, following a trend of gradual decrease in the amount of bacteria quantified on a daily interval. Antibiotics (local and systemic) were maintained for another two weeks with the concentration ramped up for local supradural treatment (from 6.6% to 16.4%) (until 64 days post-implant). In the future, a more viscous fluid (e.g., vacuum grease) may be used inside the guide tube to help prevent leakage. However, the higher viscosity fluid may also damage the fragile CF during the insertion procedure. Testing will need to be performed *in vitro* to ensure that these processes do not compromise the function of the CF sensors for neurochemical recording, and to ensure that the seal is strengthened to prevent leakage.

### 3.3. Bone Growth

Prominent bone growth was observed in both monkeys within and around the chamber interfaces (**Fig. 7**). Bone expansion towards the inner walls of the chamber was noted when the chamber cavities were exposed to install the top plate after an intermediate plate in the third procedure (126 days post-implant of the baseplate in monkey T) or to perform the craniotomy and install the top plate in the second procedure (346 days post-implant of baseplate in monkey J). White, semi-rigid, membrane-like surfaces were also observed overlying and attached to the bone in both monkeys, which were unlike normal granulation tissue that is usually found to grow above bone. Thickening of bone was also noticed at the outer perimeter of the chamber (i.e., baseplate) and skin margin. Nevertheless, these results concern external surfaces, outside of the direct interface between the bone and the PEKK baseplate. Post-terminal histology will be performed in future work to directly characterize the amount of bone expansion into the PEKK baseplate. It should also be noted that such bone growth is possible with other chamber materials, and even with the use of acrylic based cement. Future work to compare the osseointegration across different types of chamber configurations will be useful to optimize chamber design based on specific needs and/or applications.

**Figure 7.**
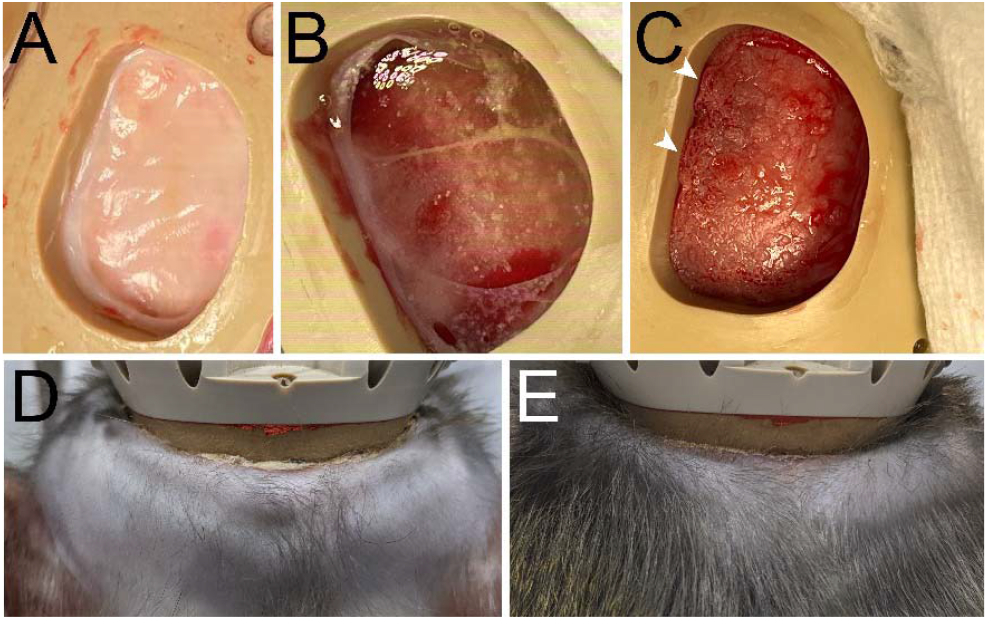
Bone growth and chamber margins. **A.** Solid membrane-like surface observed attached to bone in the chamber cavity (346 days post-baseplate implantation, monkey J). **B**. Membrane-like tissue floating and attached to bone in the perimeters of the chamber cavity (126 days post-baseplate implantation, monkey T). **C**. Bone growth observed, especially at the edges of the chamber (arrowheads) (monkey T). **D**. Posterior margin of the chamber 515 days post-baseplate implantation. **E**. Same as (**D**), 703 days post-baseplate implantation.

### 3.4. Chamber Margins

The margins of the chamber in both monkeys remained relatively healthy and intact (**Fig. 7D** and **E**). Most monkeys with chamber implants require frequent (daily to monthly) cleaning and disinfection of the margin surrounding the chamber due to skin recession, and infection, such as in the form of purulent discharge. The use of cement around the chamber usually amplifies these issues^13,14^. A small amount of purulunt exudate was observed in monkey T’s posterior margin, where the skin had shown recession. This recession was minimal and was mostly observed in an area where a wound had formed due to irritation above sharp edges of the baseplate, first appearing approximately a month after the first baseplate implantation procedure. The margins were cleaned with diluted betadine and saline every 1 – 2 weeks in monkey T. Monkey J did not undergo any margin cleaning procedures. In future procedures, the edges of the baseplate may be filleted to reduce irritation and ensure that the skin remains intact and healthy around the implanted baseplate.

### 3.5. Functional Yield for Chemical and Electrical Neural Activity Measurements

Recordings began 2 – 3 months after the implantation procedure to determine and select functional electrodes for measuring behaviorally relevant signals in the awake monkey. 53% (21/40) and 25% (13/51) of the implanted sCFE’s retained functionality (background current > 500 nA and noise < 0.05 nA) in monkeys T and J, respectively. FSCV background current was acquired during the electrode implantation procedure and microdrive lowering allowing us to identify electrodes that broke during the process of insertion. Background current is representative of the sensor sensitivity, whereas noise levels indicate the effective limit of detection (LOD) of the sensor and threshold values for these have been previously empirically determined to capture physiological (nanomolar) dopamine concentration changes *in vivo*^4,29,30^. All implanted electrodes were fully functional, as tested *in vitro*, prior to insertion. 73% (29/40) and 39% (20/51) of the implanted sCFE’s retained viable background current > 500 nA, and of these several (28% and 35% in monkeys T and J, respectively) fell short of full functionality with noise levels > 0.05 nA. It is not clear why noise is augmented for some electrodes after brain insertion, but it could be related to perforation of the insulating silica tube or parylene coating or an unstable electrical contact between the embedded CF and tungsten wire, or the tungsten and the Mill-Max pin. The yield is similar to prior work where insertions were made in a standard chamber, which is expected since the process of lowering the electrodes is the same^2^. We suspect that our low electrode insertion yield is likely due to buckling and fracture of the fragile CF during manual lowering into the brain. The fragile and thin (7 µm) CF tips are known to be susceptible to breakage during brain insertion^3^. We followed a procedure established previously for lowering of conventional electrical recording Pt/Ir and tungsten microelectrodes^11^, which we have found to rarely break during manual insertion processes. However, the speed of insertion is highly variable during manual insertion procedures and likely a significant factor in these failures. CF electrode tips also fractured upon encountering the dura mater, before they had been lowered in the brain. This occurred when the electrode was unintentionally lowered before the protective guide tube had penetrated and created a channel through the dura mater. In other cases, we suspect that the CF tips encountered other rigid surfaces or materials in the targeted trajectory, such as the meningeal layers within sulci and/or the ventricles, and/or blood vessels^31^. This is further confirmed by the observation that several electrodes (5 in monkey T, and 1 in monkey J) failed after being lowered < 1 mm on the microdrive to target new sites, months after the implant procedure.

On the other hand, the mCFE’s retained a lower functional yield of 21.4% (3/14) and 14.2% (2/14) in T and J, respectively. The thin parylene-coated CF, approximately 10 µm in diameter, extends a greater length (5 – 15 mm) from the rigid silica shaft compared to the sCFE (0.1 – 0.3 mm). As a result, the fiber is subjected to greater lateral bending forces during insertion, making it more prone to buckling. Preliminary brain phantom (i.e., 0.6% agar) simulations show that slow insertion speeds (~ 0.1 – 0.2 mm / s) reduced the probability of electrode buckling. In future studies, we plan to incorporate the use of a motorized linear-actuator to lower electrode-microdrive assemblies into the brain at a fixed velocity to reduce the risk of electrode breakage. Customized jigs will need to be manufactured to hold individual elements of the electrode-microdrive assemblies, including loosely attached guide tubes.

Some electrodes were found to have broken spontaneously during the 2 – 3 month recovery period and were likely due to an electrical disconnect between the terminal end of the electrode and Mill-Max pin that crimps it to connect to the EIB. The large movements, dynamic loads, vibrations generated by unrestrained animals in their homecage likely caused incremental stress and fatigue of this mechanical junction over time. This was especially significant in monkey J, which we suspect is due to insufficient use of varnish and liquid insulating tape at the EIB junctions, and/or the larger body mass and strength of this animal. Another challenge was electrodes getting stuck in the guide tube, therefore becoming immovable with the microdrive. This occurred in 5 CFEs in monkey T.

Of the functional CFEs, 63% (15/24) and 53% (8/15) in monkeys T and J, respectively, displayed visible dopamine signals (i.e., current changes along dopamine redox potentials), and were successfully applied in measurements during task performance. Despite the functional loss of multiple electrodes, measurements of dopamine, spike, and LFP signals were acquired successfully in the remaining sensors (**Figs. 8** and **9**). Recordings are ongoing and demonstrate the ability to monitor, chronically, molecular and electrical activity from the brain over a year.

**Figure 8.**
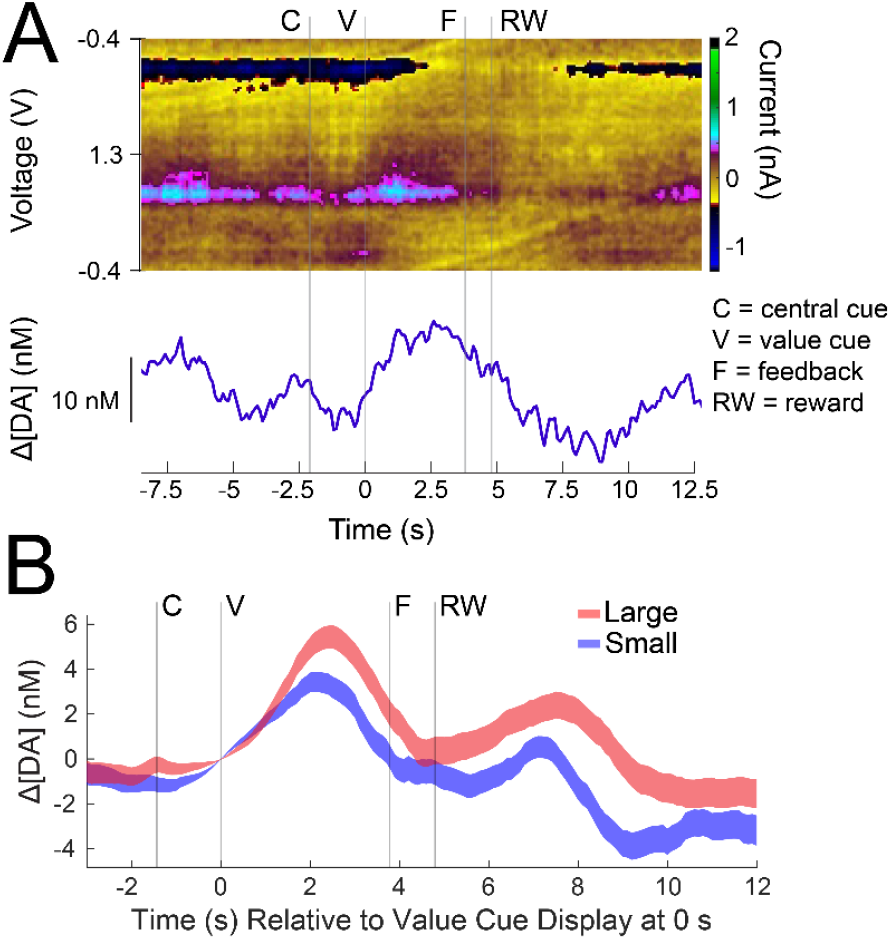
Measurements of dopamine from a chamber-implanted sCFE (site c8ds) in monkey T, 497 days post-electrode implant. **A.** Single trial measurements of dopamine concentration changes (Δ[DA]) as a function of time relative to value cue display (V) at 0 s (trial 133 in the session). The color plot is displayed on the top panel and is used to visualize selective dopamine redox current (color scale) that occurs around –0.2 and 0.6 V. The trace below it is the PCA extracted Δ[DA]. Δ[DA] fluctuations are especially visible after the value cue display. B. Trial averaged Δ[DA] aligned to V at 0 s for large and small reward trials, showing increased dopamine release for large in comparison to small reward trials, especially between the value cue display and reward (RW) event. Background subtraction occurs for all trials at 0 s to show changes relative to the value cue display event.

**Figure 9.**
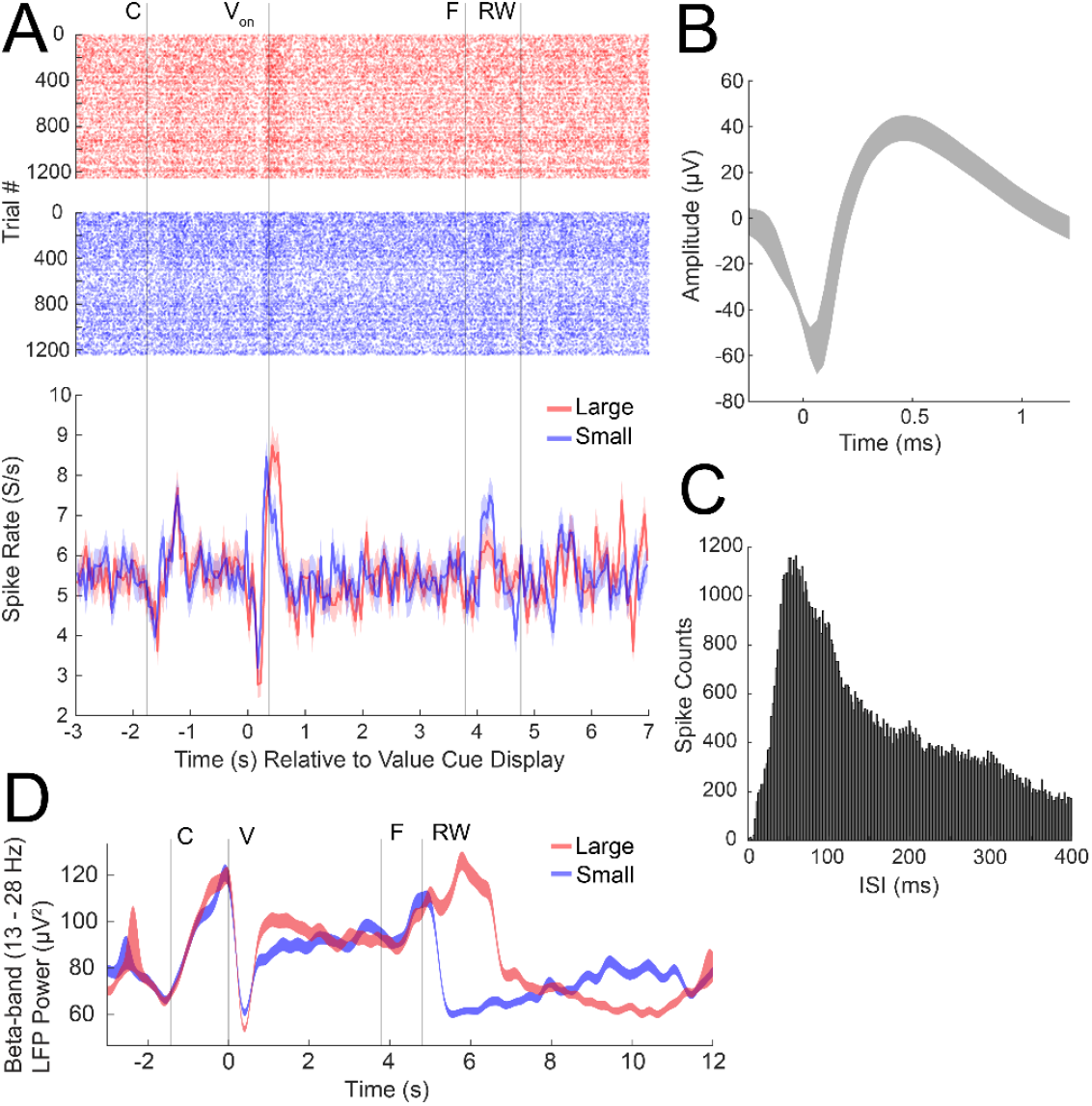
Measurements of spikes and beta-band LFPs from implanted CFEs in monkey T, 497 days post-electrode implant. **A.** (Top panel) Raster plot of spikes detected on each trial (y-axis) as a function of time relative to value cue display (V) at 0 s (site c5d). (Bottom panel) Peristimulus time histogram (PSTH) showing average spike firing rate (y-axis) for the large (red) and small (blue) reward conditions relative to the same value cue display event. V_on_ indicates average time at which fixation starts on the value cue following cue display. Event labels are the same as **Fig. 8. B**. Average spike waveform +/-standard deviation. **C**. Interspike interval (ISI) histogram showing the distribution of the timing between spikes detected. Bin widths are 1 ms. **D**. Trial averaged beta-band LFP power showing increased suppression during the brief time window immediately following the cue (V) for large as compared to small reward cues (site c7b-c5c).

## Author contributions

H.N.S., J.C., U.A., and R.M. designed and performed research, and analyzed data. H.N.S., J.C., U.A., R.M., and R.S. created implanted components. T.T., B.G., J.K.O., and D.J.S. contributed analytic tools. H.N.S., J.C., and U.A. wrote the paper.

## Acknowledgment

The authors thank Ms. Rebecca Marflak, Mr. Colum Murphy, Dr. Esta Abelev, Dr. Daniel Lamont, Ms. Erica N. Griffith, Ms. Baylie S. Leveto, Mr. Kevin Thiel, the laboratory of Dr. Tamir Ibrahim and the 7T Bioengineering Research Program (7TBRP), the Nanofabrication and Characterization Core Facility (RRID:SCR_025124), and animal care staff at the Systems Neuroscience Center Animal Resource Laboratory (SNARL) at the University of Pittsburgh (Pitt).

## Funding sources

NIH/NINDS (R00 NS107639 to H.N.S.), NIH/NINDS (R01 NS13304 to H.N.S.), the Michael J. Fox Foundation for Parkinson’s Research (MJFF) and the Aligning Science Across Parkinson’s (ASAP) initiative (ASAP-020519 to H.N.S.), and the National Defense Science and Engineering Graduate (NDSEG) Fellowship Program, sponsored by the Air Force Research Laboratory (AFRL), the Office of Naval Research (ONR), and the Army Research Office (ARO) (FA9550-21-F-0003 to U.A.).

## Declaration of interest

Baldwin Goodell is an owner of Gray Matter Research. The other authors declare no competing interests.

## Notes

### Competing Interest Statement

Charles Gray and Baldwin Goodell are owners of Gray Matter Research.

## References

1. Amjad, U., Choi, J., Gibson, D. J., Murray, R., Graybiel, A. M. & Schwerdt, H. N. Synchronous Measurements of Extracellular Action Potentials and Neurochemical Activity with Carbon Fiber Electrodes in Nonhuman Primates. eNeuro (2024). doi:10.1523/ENEURO.0001-24.2024

2. Schwerdt, H. N., Amemori, K., Gibson, D. J., Stanwicks, L. L., Yoshida, T., Bichot, N. P., Amemori, S., Desimone, R., Langer, R., Cima, M. J. & Graybiel, A. M. Dopamine and beta-band oscillations differentially link to striatal value and motor control. Science Advances 6, eabb9226 (2020).

3. Schluter, E. W., Mitz, A. R., Cheer, J. F. & Averbeck, B. B. Real-Time Dopamine Measurement in Awake Monkeys. PLOS ONE 9, e98692 (2014).

4. Schwerdt, H. N., Shimazu, H., Amemori, K., Amemori, S., Tierney, P. L., Gibson, D. J., Hong, S., Yoshida, T., Langer, R., Cima, M. J. & Graybiel, A. M. Long-term dopamine neurochemical monitoring in primates. Proceedings of the National Academy of Sciences 114, 13260–13265 (2017).

5. Yoshimi, K., Kumada, S., Weitemier, A., Jo, T. & Inoue, M. Reward-Induced Phasic Dopamine Release in the Monkey Ventral Striatum and Putamen. PLoS One 10, e0130443 (2015).

6. Boulet, S., Mounayar, S., Poupard, A., Bertrand, A., Jan, C., Pessiglione, M., Hirsch, E. C., Feuerstein, C., François, C., Féger, J., Savasta, M. & Tremblay, L. Behavioral Recovery in MPTP-Treated Monkeys: Neurochemical Mechanisms Studied by Intrastriatal Microdialysis. J. Neurosci. 28, 9575–9584 (2008).

7. Yagishita, S., Hayashi-Takagi, A., Ellis-Davies, G. C. R., Urakubo, H., Ishii, S. & Kasai, H. A critical time window for dopamine actions on the structural plasticity of dendritic spines. Science 345, 1616–1620 (2014).

8. Shindou, T., Shindou, M., Watanabe, S. & Wickens, J. A silent eligibility trace enables dopamine-dependent synaptic plasticity for reinforcement learning in the mouse striatum. European Journal of Neuroscience 49, 726–736 (2019).

9. Reynolds, J. N. J., Hyland, B. I. & Wickens, J. R. A cellular mechanism of reward-related learning. Nature 413, 67–70 (2001).

10. Dotson, N. M., Hoffman, S. J., Goodell, B. & Gray, C. M. A Large-Scale Semi-Chronic Microdrive Recording System for Non-Human Primates. Neuron 96, 769–782.e2 (2017).

11. Feingold, J., Desrochers, T. M., Fujii, N., Harlan, R., Tierney, P. L., Shimazu, H., Amemori, K.-I. & Graybiel, A. M. A system for recording neural activity chronically and simultaneously from multiple cortical and subcortical regions in nonhuman primates. J Neurophysiol 107, 1979–1995 (2012).

12. Mulliken, G. H., Bichot, N. P., Ghadooshahy, A., Sharma, J., Kornblith, S., Philcock, M. & Desimone, R. Custom-fit radiolucent cranial implants for neurophysiological recording and stimulation. Journal of Neuroscience Methods 241, 146–154 (2015).

13. Adams, D. L., Economides, J. R., Jocson, C. M., Parker, J. M. & Horton, J. C. A watertight acrylic-free titanium recording chamber for electrophysiology in behaving monkeys. Journal of Neurophysiology 106, 1581–1590 (2011).

14. Lanz, F., Lanz, X., Scherly, A., Moret, V., Gaillard, A., Gruner, P., Hoogewoud, H.-M., Belhaj-Saif, A., Loquet, G. & Rouiller, E. M. Refined methodology for implantation of a head fixation device and chronic recording chambers in non-human primates. Journal of Neuroscience Methods 219, 262–270 (2013).

15. Overton, J. A., Cooke, D. F., Goldring, A. B., Lucero, S. A., Weatherford, C. & Recanzone, G. H. Improved methods for acrylic-free implants in nonhuman primates for neuroscience research. Journal of Neurophysiology 118, 3252–3270 (2017).

16. Gray, C. M., Goodell, B. & Lear, A. Multichannel micromanipulator and chamber system for recording multineuronal activity in alert, non-human primates. J Neurophysiol 98, 527–536 (2007).

17. Watanabe, K., Kadohisa, M., Kusunoki, M., Buckley, M. J. & Duncan, J. Cycles of goal silencing and reactivation underlie complex problem-solving in primate frontal and parietal cortex. Nat Commun 14, 5054 (2023).

18. Amemori, S., Graybiel, A. M. & Amemori, K. Cingulate microstimulation induces negative decision-making via reduced top-down influence on primate fronto-cingulo-striatal network. Nat Commun 15, 4201 (2024).

19. Zhang, K., Bromberg-Martin, E. S., Sogukpinar, F., Kocher, K. & Monosov, I. E. Surprise and recency in novelty detection in the primate brain. Current Biology 32, 2160–2173.e6 (2022).

20. Wang, M., Bhardwaj, G. & Webster, T. J. Antibacterial properties of PEKK for orthopedic applications. International Journal of Nanomedicine 12, 6471–6476 (2017).

21. Chang, C.-C. & Merritt, K. Microbial adherence on poly(methyl methacrylate) (PMMA) surfaces. Journal of Biomedical Materials Research 26, 197–207 (1992).

22. Yuan, B., Cheng, Q., Zhao, R., Zhu, X., Yang, X., Yang, X., Zhang, K., Song, Y. & Zhang, X. Comparison of osteointegration property between PEKK and PEEK: Effects of surface structure and chemistry. Biomaterials 170, 116–126 (2018).

23. Hong, S.-O., Pyo, J.-Y., On, S.-W., Seo, J.-Y. & Choi, J.-Y. The Biocompatibility and the Effect of Titanium and PEKK on the Osseointegration of Customized Facial Implants. Materials 17, 4435 (2024).

24. Cheng, B. C., Jaffee, S., Averick, S., Swink, I., Horvath, S. & Zhukauskas, R. A comparative study of three biomaterials in an ovine bone defect model. The Spine Journal 20, 457–464 (2020).

25. Schwitalla, A. & Müller, W.-D. PEEK Dental Implants: A Review of the Literature. Journal of Oral Implantology 39, 743–749 (2013).

26. Liang, L., Zimmermann Rollin, I., Alikaya, A., Ho, J. C., Santini, T., Bostan, A. C., Schwerdt, H. N., Stauffer, W. R., Ibrahim, T. S., Pirondini, E. & Schaeffer, D. J. An open-source MRI compatible frame for multimodal presurgical mapping in macaque and capuchin monkeys. Journal of Neuroscience Methods 407, 110133 (2024).

27. Santini, T., Wood, S., Krishnamurthy, N., Martins, T., Aizenstein, H. J. & Ibrahim, T. S. Improved 7 Tesla transmit field homogeneity with reduced electromagnetic power deposition using coupled Tic Tac Toe antennas. Sci Rep 11, 3370 (2021).

28. Keithley, R. B. & Wightman, R. M. Assessing principal component regression prediction of neurochemicals detected with fast-scan cyclic voltammetry. ACS Chem Neurosci 2, 514–525 (2011).

29. Schwerdt, H. N., Zhang, E., Kim, M. J., Yoshida, T., Stanwicks, L., Amemori, S., Dagdeviren, H. E., Langer, R., Cima, M. J. & Graybiel, A. M. Cellular-scale probes enable stable chronic subsecond monitoring of dopamine neurochemicals in a rodent model. Commun Biol 1, 1–11 (2018).

30. Schwerdt, H. N., Kim, M. J., Amemori, S., Homma, D., Yoshida, T., Shimazu, H., Yerramreddy, H., Karasan, E., Langer, R., Graybiel, A. M. & Cima, M. J. Subcellular probes for neurochemical recording from multiple brain sites. Lab Chip 17, 1104–1115 (2017).

31. Reiter, N., Paulsen, F. & Budday, S. Mechanisms of mechanical load transfer through brain tissue. Sci Rep 13, 8703 (2023).

